# Comparative lifespan trajectories of brain energy metabolism in human and macaque

**DOI:** 10.64898/2026.04.16.719061

**Authors:** Moohebat Pourmajidian, Bratislav Misic, Alain Dagher

## Abstract

The brain’s extraordinary energy demands are met by a suite of metabolic pathways that selectively appropriate glucose across development, yet how this metabolic strategy changes across the lifespan and whether it is conserved across species remains incompletely understood. We previously mapped five core energy metabolism pathways across the human cortex and lifespan, revealing a fundamental dichotomy between anabolic and energy-producing pathway expression: the pentose phosphate pathway, which contributes to biomass, peaks prenatally, while glycolysis, the TCA cycle, and oxidative phosphorylation rise postnatally. Here, we extend the analysis to the rhesus macaque and show that the prenatal-to-postnatal transition is similar across the two species. Using the MitoCarta3.0 annotation framework, we further identify two consistent mitochondria-specific programs: a progressive decline in mitochondrial genome maintenance pathways, and a postnatal rise in mitochondrial energy production and substrate utilization, replicated across human and macaque. Extending to mouse, rat, and chicken, we find this shifting metabolic strategy is a conserved feature of vertebrate brain development. Finally, we map the cortical expression of mitochondria-localized pathway genes in the adult human cortex, finding that the anabolic-oxidative dichotomy follows in the mature brain. Together, these findings provide a comparative transcriptomic framework for studying brain energy metabolism across the lifespan.

## INTRODUCTION

Energy metabolism is a fundamental process in cellular organisms. It provides fuel for cellular function and homeostasis and substrates for tissue biosynthesis and repair (Bauernfeind et al. 2014, McKenna et al. 2019). In the brain, these demands are exceptionally high: neuronal signaling at the resting state alone imposes a high energetic cost, and the reactive oxygen species (ROS) generated during mitochondrial respiration render the brain particularly vulnerable to oxidative damage (Attwell and Laughlin 2001, Lin and Beal 2006, Magistretti and Allaman 2015). Energy metabolism pathways utilize glucose and lactate to produce adenosine triphosphate (ATP), the energy currency for all functions within the cell. However, by producing intermediates for the synthesis of fatty acids, nucleotides and amino acids, and molecules involved in redox buffering, these pathways also contribute to tissue building and oxidative defense (Camandola and Mattson 2017, Magistretti and Allaman 2015).

Within the cell, glucose is converted to pyruvate via glycolysis, producing two ATP molecules. Pyruvate is then transported into the mitochondria, where it is converted to acetyl-CoA and enters the tricarboxylic acid (TCA) cycle, producing high-energy electron carriers that drive ATP synthesis through the mitochondrial electron transport chain (ETC) and oxidative phosphorylation (OXPHOS). Furthermore, lactate dehydrogenase catalyzes the interconversion of pyruvate and lactate, allowing lactate to serve as an oxidative fuel in the TCA cycle (Magistretti 2006, Magistretti and Allaman 2018). Acetyl-coA and other TCA intermediates also play integral roles in fatty acid and amino acid synthesis. Parallel to its oxidative fate, glucose can enter the pentose phosphate pathway (PPP) to produce 5-carbon sugars used in the synthesis of nucleotides and amino acids. The PPP also produces nicotinamide adenine-dinucleotide phosphate (NADPH), an important co-factor in lipid synthesis and oxidative defense via the glutathione cycle (Camandola and Mattson 2017, Magistretti and Allaman 2015).

Brain energy metabolism changes throughout the lifespan. An especially dynamic period is the transition from prenatal to postnatal stages, when metabolic demands shift in response to rapidly changing developmental and nutritional needs. (Bauernfeind and Babbitt 2014, McKenna et al. 2019). In the prenatal brain, anabolic demands are high, supporting neural proliferation and tissue growth (Baquer et al. 1988, Brekke et al. 2015, Morken et al. 2014). As the brain matures postnatally, a shift toward oxidative energy production occurs. PET imaging studies have shown that cerebral glucose uptake and oxygen consumption increase after birth and peak in childhood (Chugani 1998, Kuzawa et al. 2014, McKenna et al. 2012), and subsequently decline through adolescence and into adulthood (Kennedy and Sokoloff 1957, Takahashi et al. 1999). In parallel, aerobic glycolysis, a measure of non-oxidative glucose utilization, is also reported to progressively decline in the aging brain (Goyal et al. 2014, Vaishnavi et al. 2010).

We recently characterized energy metabolism programs at the pathway level in the human brain, mapping the spatial and temporal pattern of gene expression for five core energy metabolism pathways, namely glycolysis, PPP, TCA cycle, OXPHOS, and lactate, using spatially-comprehensive gene expression atlases (Pourmajidian et al. 2026). This approach revealed consistent spatial heterogeneity, with motor cortices showing high glycolysis and OXPHOS expression, and primary sensory regions enriched for PPP-related genes. Across development, PPP expression was most pronounced prenatally and declined after birth, while pathways mainly involved in oxidative metabolism of glucose and ATP production showed a postnatal increase and peaked in childhood. This transcriptomic framework offers pathway-level specificity that is inaccessible to current neuroimaging modalities, which provide measures of glucose uptake without resolving the downstream metabolic fate of glucose. Yet, a fundamental question remains open: are these temporal patterns unique to the human brain, or do they reflect shared metabolic programs across the evolutionary history?

Core metabolic pathways, such as glycolysis, PPP and TCA, show remarkable conservation not only in the high sequence similarity of their constituent genes and proteins, but also in their reaction sequence, preserved across billions of years of diverging evolution (Peregrín-Alvarez et al. 2009, Ralser et al. 2021). On the other hand, several genes encoding components of the mitochondrial electron transport chain have been subject to positive selection in the anthropoid primate lineage, with accelerated rates of non-synonymous substitution in subunits of complexes III and IV, suggesting that remodeling of the OXPHOS system accompanied the expansion of the energetically expensive neocortex (Grossman et al. 2001, 2004). Furthermore, comparative transcriptomic studies have consistently identified up-regulation of genes involved in glucose metabolism and mitochondrial oxidative phosphorylation in the human brain relative to non-human primates (Bauernfeind et al. 2015, Cáceres et al. 2003, Khaitovich et al. 2006b, Uddin et al. 2008, 2004). Metabolomic analyses further show that energy metabolism in the human prefrontal cortex has undergone accelerated evolutionary change (Fu et al. 2011, Khaitovich et al. 2008). This divergence extends to the developmental dimension: compared to other primates, humans show a protracted schedule of synaptogenesis, myelination and peak glucose uptake (Alldritt et al. 2024, Somel et al. 2009), processes that impose distinct energetic demands, suggesting that the timing and allocation of metabolic programs is also subject to evolutionary divergence (Bauernfeind and Babbitt 2014, Bauernfeind et al. 2014). How these evolutionary differences manifest at the level of metabolic pathways, however, remains incompletely understood.

Here, we take a comparative approach by leveraging multiple developmental transcriptomic datasets to track the expression trajectory of key glucose and mitochondrial metabolic pathways across the human and nonhuman primate (macaque) lifespan. We report that energy pathways show a remarkable similarity between these two species, capturing the dichotomy in the appropriation of glucose toward anabolic versus ATP-producing pathways. Using the MitoCarta3.0 annotation framework, we further identify two mitochondria-specific programs: a prenatal enrichment in mitochondrial genome maintenance pathways and a postnatal rise in oxidative and metabolic pathways. Furthermore, we extend the analysis to three other mammalian and avian species and find that this metabolic dichotomy is consistently observed beyond the primate brain. Finally, we map the cortical spatial expression of mitochondrial pathways in the adult human brain, finding that the anabolic-oxidative dichotomy is extended to the mature cortex. Together, this study provides new insights into the shared metabolic programs that govern brain development and highlights the potential of transcriptomics to study cross-species dynamics across the lifespan. This work further sheds light on cross-species similarities and differences in brain energy metabolism.

## RESULTS

To characterize the developmental shifts in brain energy metabolism in the macaque, we use bulk RNA-sequencing data spanning prenatal development through adulthood from the PsychENCODE evolution dataset (Zhu et al. 2018). Briefly, gene sets for each energy pathway were obtained from the Gene Ontology biological processes (Ashburner et al. 2000) and Reactome pathways (Croft et al. 2014) as in our recent work (Pourmajidian et al. 2026). Consensus genes annotated in both databases were kept for further analysis. For a list of pathways IDs and genes see Table S1. For the mitochondria-specific pathways, gene sets were retrieved from the Mitocarta3.0 database, an inventory of genes encoding mitochondria-localized proteins in human and mouse (Calvo et al. 2016, Pagliarini et al. 2008, Rath et al. 2021). Sample by gene expression matrices were then made for each pathway and expression was averaged across all genes to produce a mean gene expression vector for each pathway. The full set of pathways and genes are available in S1 Data. The analysis is further extended to mouse, rat, and chicken using a bulk RNA-sequencing developmental dataset spanning early brain organogenesis through adulthood (Cardoso-Moreira et al. 2019), and to the adult human cortex using microarray data from the Allen Human Brain Atlas (Hawrylycz et al. 2012).

### Lifespan trajectory of key glucose metabolism pathways in human and macaque

The macaque is a foundational model organism in neuroscience, with prior work characterizing shared and divergent features of human and macaque brain across structural and functional organization, metabolism, cell types, neurotransmitter systems and developmental gene expression programs (Alldritt et al. 2024, Bakken et al. 2016, Bauernfeind and Babbitt 2014, Burt et al. 2018, Eberling et al. 1995, Jacobs et al. 1995, Kennedy et al. 1982, Luppi et al. 2025, Seidlitz et al. 2018, Vogel et al. 2024, Xu et al. 2020, Zhu et al. 2018). Yet how brain energy metabolism is appropriated across pathways throughout the macaque lifespan remains understudied. We produce gene expression trajectories for six pathways involved in brain energy metabolism: glycolysis, pentose phosphate pathway (PPP), TCA cycle, oxidative phosphorylation (OXPHOS), lactate metabolism and transport, and ketone body utilization. The median pathway expression across all samples at each age (embryonic day 60 to 11 years) is plotted (Figure 1a). To ease interpretation, we additionally summarize pathway expression across broader developmental stages (prenatal, infant, juvenile, adolescent, and adult) by pooling samples within each stage (Figure 1b). For details of ages included in each group see Table S2. Two broadly different trajectories are observed: pathways mainly involved in ATP production including glycolysis, TCA and OXPHOS show an increase in their average gene expression from the prenatal stage to infancy, while the PPP declines postnatally, reaching its lowest expression in adolescence and adulthood. OXPHOS expression subsequently declines in adolescence and adulthood. These patterns are consistent with our previous findings in the human brain (Pourmajidian et al. 2026). The decline in OXPHOS expression could also parallel previous reports of age-associated reductions in cerebral glucose uptake and blood flow in PET imaging studies of young and aged macaques (Cross et al. 2000, Eberling et al. 1997, Noda et al. 2002).

**Figure 1.**
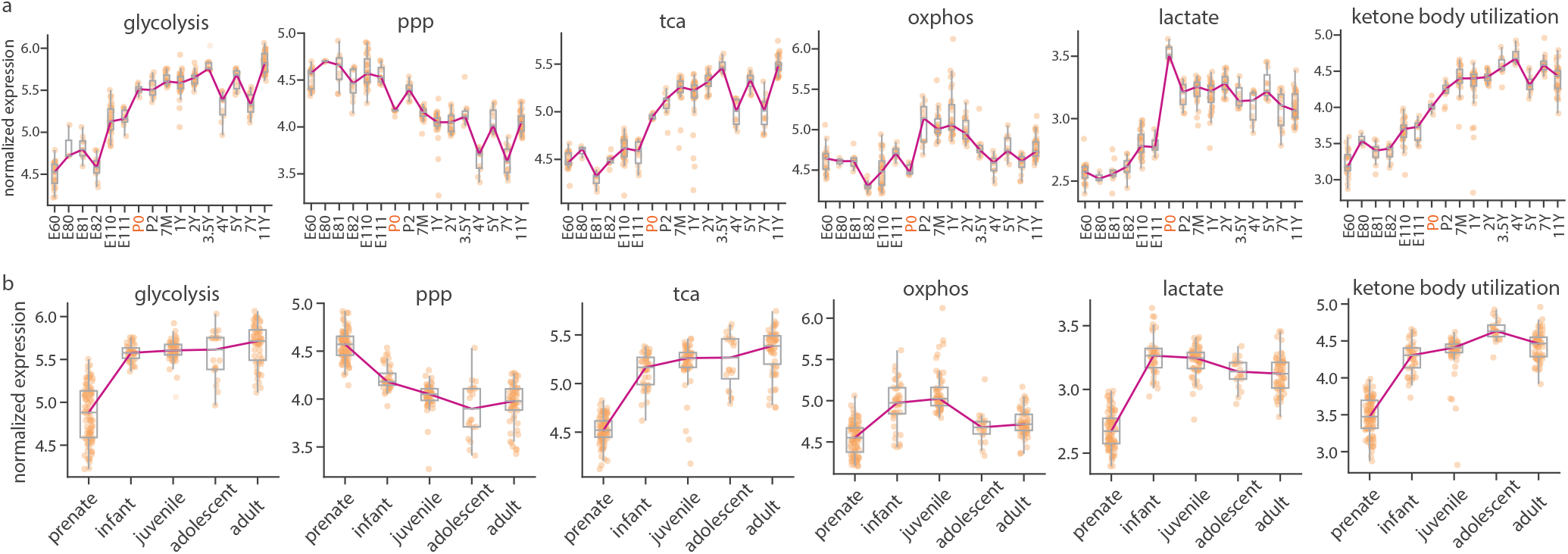
Lifespan trajectory of energy metabolism pathways in the rhesus macaque. Developmental trajectories of energy metabolism pathways were produced using the Psychencode evolution RNA-sequencing dataset http://evolution.psychencode.org. For each energy pathway, mean expression was calculated across all genes for each sample. Analysis only included cortical regions. (a) Median expression across all samples at each age. Dots represent individual samples at each age. (b) Median expression across all samples at each age group. For details of ages included in each group see Table S2. The y-axis represents normalized log_2_(RPKM) expression values (*See Methods*). Pathway gene sets used to create these traejctories can be found in S1 Data. ppp, pentose phosphate pathway; tca, tricarboxylic acid cycle; oxphos, oxidative phosphorylation; lactate, lactate metabolism and transport.

To directly compare developmental trajectories in human and macaque, ages were converted to post-conception days (PCD) and log_10_-transformed to enable a common developmental axis (Zhu et al. 2018). Given the limited number of samples and coverage per time-point, comparisons are restricted to the broad shape of the trajectories rather than quantitative differences in timing (Figure 2). Across all six energy pathways, human and macaque show similar expression trajectories (Figure 2a).

**Figure 2.**
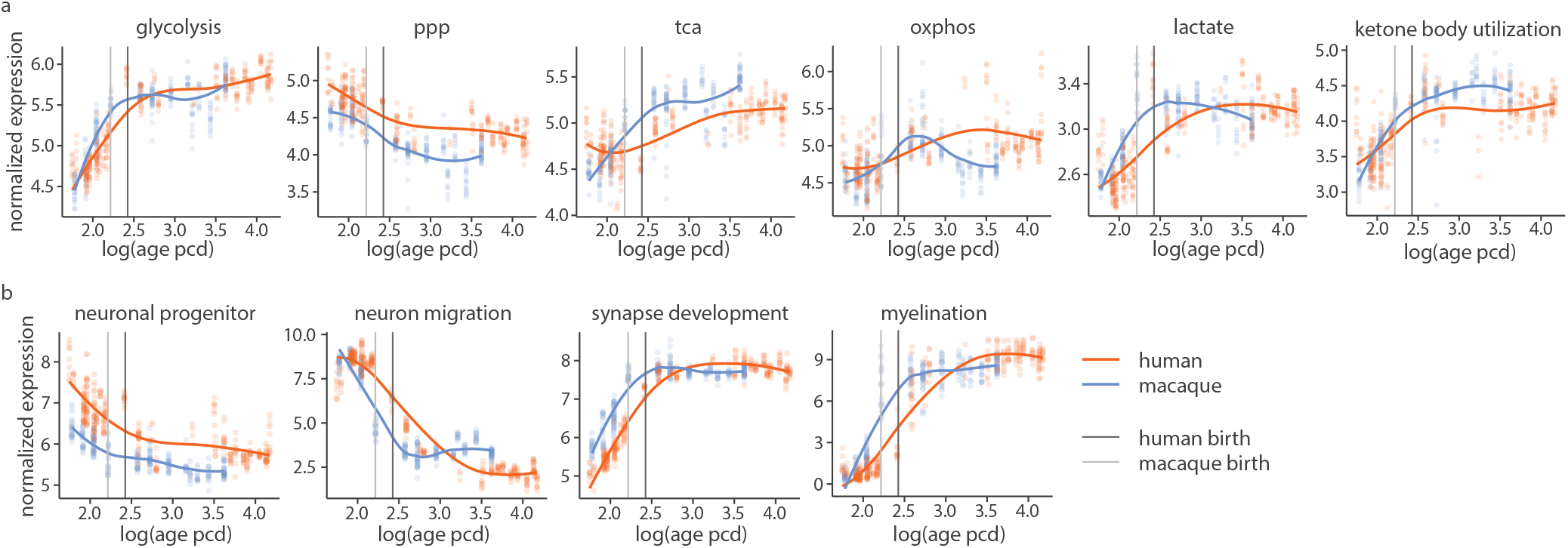
Lifespan trajectories of energy metabolism pathways across the human and macaque. Using both the human and macaque samples in the PsychENCODE evolution dataset, we fit lifespan trajectories using the non-parametric locally estimated scatterplot smoothing (LOESS) method. For the number of samples at each spatiotemporal point see S8. (a) For each energy pathway, mean expression was calculated across all genes for each sample. (b) Using previously curated gene sets (Kang et al. 2011, Li et al. 2018), mean expression of marker genes for each developmental process was calculated for each sample. The x-axis represent log_10_transformed age in post conception days (PCD). Dots represent individual cortical samples at each age. The y-axis represents normalized log_2_(RPKM) expression values (*See Methods*). Pathway gene sets used to create these traejctories can be found in S1 Data. ppp, pentose phosphate pathway; tca, tricarboxylic acid cycle; oxphos, oxidative phosphorylation; lactate, lactate metabolism and transport.

To contextualize these patterns within the broader neurodevelopmental landscape, we additionally plot expression trajectories for marker gene sets of neural progenitor cells, neuronal migration, synapse development, and myelination (Figure 2b). Both species show concordant postnatal declines in neural progenitor and neuronal migration markers and concordant rises in synaptic development and myelination programs. This parallels the shift from anabolic PPP metabolism in the prenate to oxidative metabolism postnatally, consistent with the role of the PPP in supporting tissue generation and neurogenesis, and the subsequent reliance on glycolysis and oxidative metabolism to meet the energetic demands of synaptic maturation.

### Mitochondrial function across the human and macaque lifespan

Mitochondrial function arises from the coordinated expression of nuclear- and mitochondrially-encoded proteins. To systematically characterize the developmental organization of mitochondrial function, we examined the lifespan expression trajectories of pathways annotated in MitoCarta3.0, a curated inventory of mitochondria-localized proteins organized into 149 functional pathways across a three-tier hierarchy (Rath et al. 2021). The first level contains seven broad mitochondrial processes: mitochondrial central dogma, protein import, sorting and homeostasis, oxidative phosphorylation, metabolism, small molecule transport, mitochondrial dynamics and surveillance, and signaling, while the second and third levels provide progressively finer functional subdivisions within each process. We focused on a subset of pathways at the second and third hierarchy levels relevant to energy metabolism and which showed consistent trajectories across independent platforms and summary measures (see Methods; Figures S1, S2).

Two broad patterns emerge consistently from the mitochondria-specific pathways (Figure 3). First, pathways associated with mitochondrial genome maintenance, including mtDNA replication, mtDNA nucleoid, mtDNA repair and mitochondrial transcription, show elevated expression prenatally that declines after birth, in both species. This prenatal enrichment may be consistent with the need to maintain and replicate the mitochondrial pool during rapid division of neural progenitor cells in early brain development. The coordinated postnatal decline of these pathways across both human and macaque suggests that this transition may be a conserved feature of primate brain maturation.

**Figure 3.**
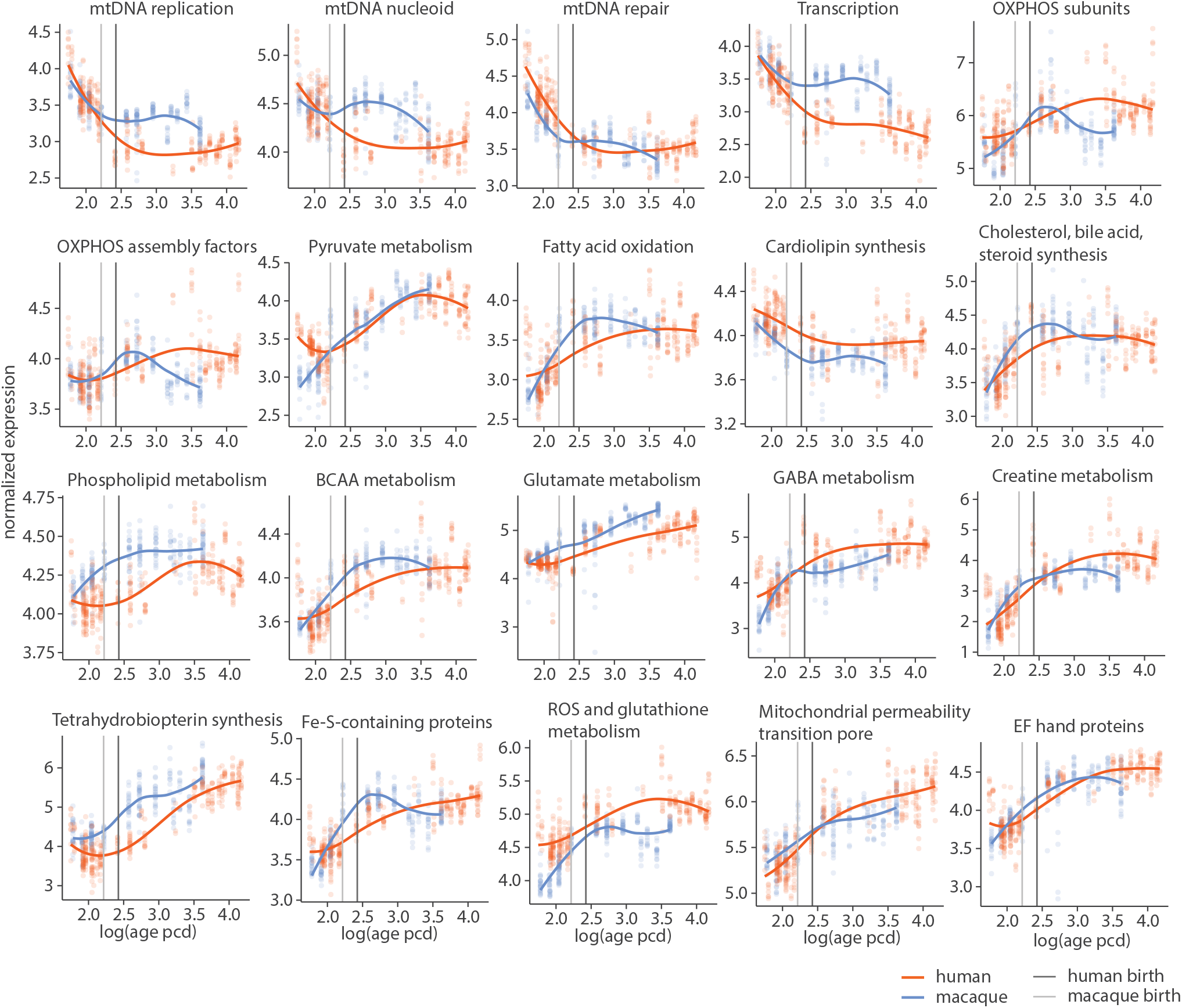
Lifespan trajectories of Mitocarta3.0 pathways in human and macaque. The human and macaque sample-by-gene expression matrices were filtered using pathway gene sets from the Mitocarta3.0 database (Rath et al. 2021). For each energy pathway, mean expression was calculated across all genes for each sample. Only cortical samples were included. Lifespan trajectories were modeled using the non-parametric locally estimated scatterplot smoothing (LOESS) method. The y-axis represents normalized log_2_(RPKM) expression values (*See Methods*). The x-axis represent log_10_transformed age in post conception days (PCD). Dots represent individual cortical samples at each age. MitoCarta3.0 pathway genes sets used to create these trajectories can be found in S1 Data. ppp, pentose phosphate pathway; tca, tricarboxylic acid cycle; oxphos, oxidative phosphorylation; lactate, lactate metabolism and transport.

Second, pathways associated with mitochondrial energy production show a consistent postnatal rise in both species. These include the expression of genes in the OXPHOS machinery, pyruvate metabolism, fatty acid oxidation, creatine metabolism, ROS and glutathione metabolism. The postnatal rise in OXPHOS complex expression is consistent with the increased mitochondrial respiratory capacity required to sustain synaptic energy demands during the critical period of synaptic development (Harris et al. 2012, Kuzawa et al. 2014, Zheng et al. 2016), and aligns with a previously reported conserved postnatal elevation of mitonuclear OXPHOS gene expression across vertebrates in the same dataset (Medini and Mishmar 2025). The concurrent rise in ROS and glutathione metabolism pathways may reflects an adaptive antioxidant response to the increased ROS load associated with mitochondrial respiration (Murphy 2009, Turrens 2003). We also see a postnatal increase in expression of genes involved in lipid metabolism, including cholesterol, bile acid, and steroid synthesis, and phospholipid metabolism. This parallels the progression of myelination, linking postnatal mitochondrial metabolic activity to the structural maturation of white matter. Inspection of the trajectories suggests that the postnatal rise in oxidative metabolism pathways (e.g. OXPHOS, TCA, fatty acid oxidation) may be more protracted in humans relative to macaque, though the sparse sampling of the dataset and the inherent differences in developmental timing between these two species precludes confident conclusions about timing.

Taken together, our results show that the change in metabolic programs during development is broadly similar between human and macaque.

### Pre-natal and post-natal trajectory of energy pathways across the mammalian and avian species

We next extend the lifespan analyses to three other mammalian and avian species: mouse (*Mus musculus*), rat (*Rattus norvegicus*), and chicken (*Gallus gallus*), using bulk RNA-sequencing data from (Cardoso-Moreira et al. 2019) (Figure 4). As before, sample-by-gene expression matrices were built for gene sets corresponding to energy and mitochondrial pathways and the median gene expression was plotted across age. Given that each developmental time point is represented by only 3–4 samples in this dataset and the sparse coverage of post-natal stages, results are interpreted in terms of broad directional patterns rather than precise trajectory shapes.

**Figure 4.**
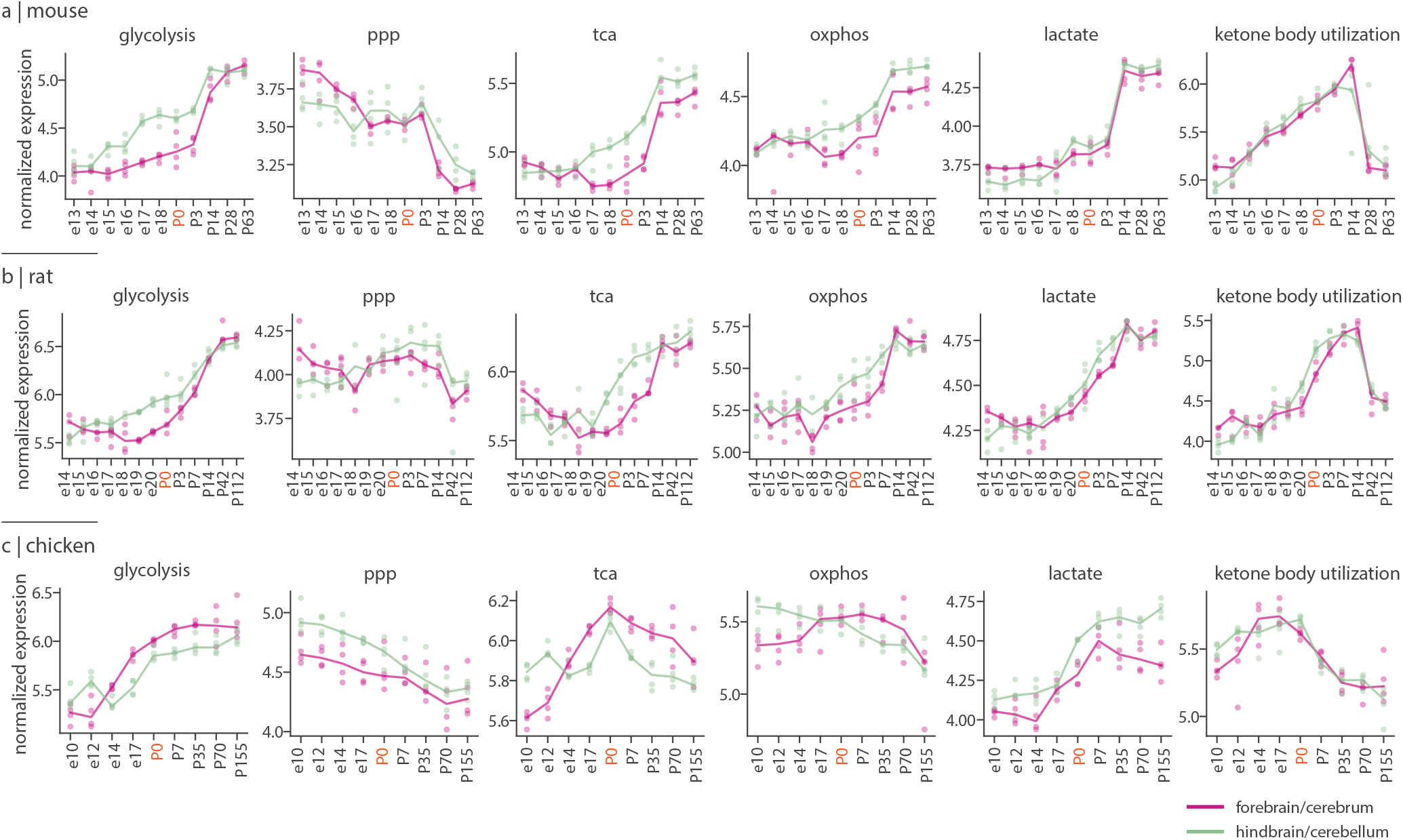
Lifespan trajectories of energy metabolism pathways across vertebrate species. Developmental trajectories of energy metabolism pathways were produced using bulk RNA-seq data from the forebrain and cerebellum of mouse (*Mus musculus*), rat (*Rattus norvegicus*), and chicken (*Gallus gallus*) (Cardoso-Moreira et al. 2019). For each energy pathway, mean expression was calculated across all genes for each sample. The median pathway expression across samples was then plotted across all ages for each species. Pink and green lines represent median gene expression in the forebrain and the cerebellum, respectively. dots represent individual samples at each age. P0 (red label) denotes birth or hatching. Dots represent individual samples. Pathway gene sets used to create these mean expression trajectories can be found in S1 Data. ppp, pentose phosphate pathway; tca, tricarboxylic acid cycle; oxphos, oxidative phosphorylation; lactate, lactate metabolism and transport.

Despite this sparse sampling, the core metabolic dichotomy identified in human and macaque is broadly recapitulated. In mouse, the pattern is most clearly apparent: PPP expression declines markedly after birth, while glycolysis, TCA, OXPHOS, and lactate all rise postnatally. In rat, glycolysis, TCA, OXPHOS, and lactate similarly rise postnatally, though PPP shows a less consistent decline compared to mouse. In chicken, glycolysis rises and PPP declines across the prenatal-to-postnatal transition, extending the core pattern to a non-mammalian vertebrate. OXPHOS, TCA and lactate also show dynamic trajectories, though their peak expression appears to happen at and around hatching with a decline afterwards. These findings are consistent with a recent analysis of mitonuclear gene expression in the same dataset, which reported a conserved upregulation of OXPHOS and TCA genes across vertebrates after birth (Medini and Mishmar 2025).

In both mouse and rat, ketone body utilization rises postnatally before declining in later postnatal stages (P28 and P42, respectively), roughly coinciding with the weaning period (approximately 3–4 weeks after birth in laboratory mice and rats (Curley et al. 2009, Kikusui and Mori 2009, Ošt’ádalová and Babický 2012). This is in alignment with the postnatal activity profile of ketone body metabolism enzymes in rat brain (Bilger and Nehlig 1992, Lockwood and Bailey 1971, Nehlig 2004) and previous autoradiographic measurements of cerebral beta-hydroxybutyrate uptake, which peak at P14 and decline by approximately 50% by P35 (Nehlig 1999, Nehlig et al. 1991).

Ketone body utilization in chicken shows a different pattern than in mammals: expression is high embryonically and declines after hatching. This likely reflects the fact that the chicken embryo relies on yolk-derived fatty acids as its primary energy source prior to hatching (Linares et al. 1993), which declines with the exhaustion of this lipid source (Nehlig et al. 1978, Nehlig and Lehr 1981). This is unlike suckling mammals whose brains are supplemented by dietary fat-derived ketones through nursing postnatally (Bilger and Nehlig 1992, Nehlig 2004).

The mitochondria-specific pathway trajectories are similarly conserved across all three species (Figure S3). In mouse and rat, pathways associated with mitochondrial genome maintenance (mtDNA replication, mtDNA nucleoid and mtDNA repair) show a progressive decline during development, beginning prenatally and continuing after birth, consistent with the patterns observed in human and macaque. Conversely, pathways associated with energy production and substrate utilization, including OXPHOS subunits, pyruvate metabolism, creatine metabolism, and neurotransmitter metabolism, show a postnatal rise in both species. In chicken, patterns are broadly similar with respect to hatching.

Taken together, these results indicate that the prenatal-to-postnatal metabolic shift is broadly conserved across mammalian and non-mammalian vertebrate species, with a prenatal anabolic state with pronounced PPP expression transitioning into a postnatal energy-producing state driven by glycolysis, TCA, and OXPHOS.

### Spatial expression of mitochondrial pathway genes in the human cortex

Finally, we map the spatial pattern of gene expression for the mitochondria-specific pathways in the adult human cortex. Microarray gene expression data from the Allen Human Brain Atlas (Hawrylycz et al. 2012) were parcellated into the Schaefer-400 cortical atlas (Schaefer et al. 2018) using the *abagen* package (Markello et al. 2021) (https://abagen.readthedocs.io/). Note that the data is from post-mortem adult donors (ages 24–57, 42.50 *±* 13.38). region-by-gene expression matrices were created for each pathway and the mean expression across all genes was calculated (Figure 5a).

**Figure 5.**
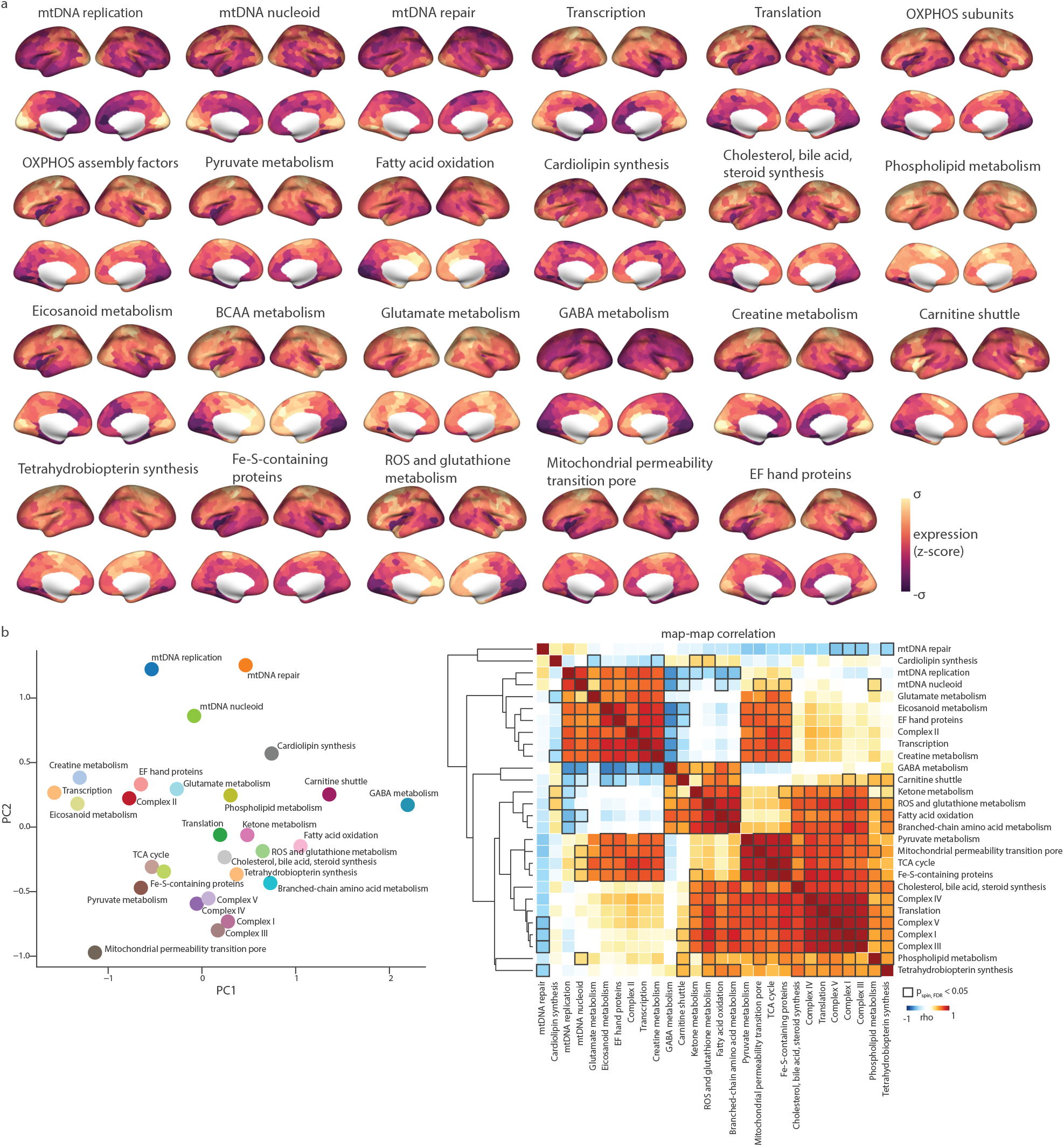
Cortical gene expression maps of mitochondria-specific pathways in the adult human brain. Microarray gene expression data from the Allen Human Brain Atlas (Hawrylycz et al. 2012) were parcellated into the Schaefer-400 cortical atlas (Schaefer et al. 2018) using the *abagen* package (Markello et al. 2021). Data are from 6 post-mortem adult donors (ages 24–57). (a) Cortical mean gene expression maps across the Schaefer-400 parcellation. For each mitochondria-specific pathway annotated in Mitocarta3.0 (Rath et al. 2021), mean expression was calculated across all pathway genes for each cortical parcel. Color bar represents mean gene expression values. (b) Clustering of MitoCarta3.0 pathway maps. Left: Clustering of the complete set of energy maps was carried out using PCA on a matrix containing all pathway mean gene expressions. Axes represent the first and second principal components. Right: Pair-wise Spearman’s correlation was calculated between the mitochondrial pathway mean gene expression maps. Dendrogram represents the hierarchical clusters. Colorbar represents Spearman’s correlation values. Bold edges indicate statistical significance tested against 1 000 spatial autocorrelation preserving nulls and after FDR correction using the Bejamini-Hochberg method for multiple comparisons. Data underlying this figure and the full set of MitoCarta3.0 pathway maps can be found in S1 Data.

Two patterns are readily observed: pathways involved in mitochondrial genome maintenance and transcription show greater expression in the posterior cortices, specifically in the visual cortex S7. This resembles the spatial pattern of the PPP previously reported and may hint at shared anabolic and mitochondrial biogenesis programs in these cortices. In contrast, pathways relating to carbohydrate and lipid metabolism and OXPHOS components show lower expression in the visual cortex and greater expression in the anterior and motor regions S7, in accord with our previous findings (Pourmajidian et al. 2026). The clustering of these maps is shown in 5b. OXPHOS complexes I, III, IV and V form a tight cluster within a broader cluster of pathways involved in fatty acid oxidation, TCA cycle activity, and ROS and glutathione metabolism. The full set of MitoCarta3.0 pathways maps are provided in S1 Data.

Collectively, the spatial organization of these pathways mirrors the temporal dichotomy identified across development, with pathways enriched prenatally and associated with anabolic and maintenance functions showing posterior enrichment in the adult brain, while pathways associated with energy production showing anterior enrichment.

### Sensitivity and replication

To ensure results are robust to analytic choices, we repeated all lifespan trajectory analyses using the first principal component (PC1) of gene expression as an alternative summary measure to mean expression. PC1 captures the axis of maximum variance across genes within each pathway gene set and is therefore less sensitive to the influence of individual gene estimations. Across all species, trajectory patterns remain consistent when using PC1 instead of mean expression (Figures S1, S4, S5). The PC1 maps of cortical gene expression for the MitoCarta3.0 pathways is also provided S6.

We additionally replicated the human mitochondrial pathway trajectories using the BrainSpan microarray dataset (Kang et al. 2011) as an independent platform. Trajectories remain broadly consistent across the two platforms, supporting the robustness of the observed developmental patterns (Figure S2).

## DISCUSSION

The advent of large-scale sequencing platforms and the growing availability of cross-species transcriptomic datasets have opened new avenues for studying the conservation and divergence of molecular programs across mammalian and non-mammalian species. Energy metabolism is among the most essential cellular functions, providing both fuel and molecular building blocks for the growth, repair, and survival of the organism. These roles are carried out by distinct pathway programs: the pentose phosphate pathway contributes to tissue biosynthesis, cellular repair and oxidative defense, while glycolysis and oxidative phosphorylation drive ATP production, with the TCA cycle bridging both functions through its anaplerotic intermediates (Camandola and Mattson 2017, Magistretti and Allaman 2015, McKenna et al. 2012). Given these essential functionalities, core energy metabolism pathways show remarkable conservation throughout evolution, both in the high sequence similarity in genes and proteins and their reaction architecture (Liska et al. 2023, Peregrín-Alvarez et al. 2009, Ralser et al. 2021). Yet, in the primate brain, and specifically humans, energy metabolism pathways have been subject to positive selection across multiple biological scales (Babbitt et al. 2011) including gene sequences, amino acid sequences, gene regulatory elements (Babbitt et al. 2010, Cáceres et al. 2003, Haygood et al. 2007, Khaitovich et al. 2006a,b), reflected also in human-specific divergence in brain metabolite profiles (Fu et al. 2011, Khaitovich et al. 2008).

Here, we take a comparative approach to construct lifespan expression trajectories for key pathways involved in brain energy metabolism and mitochondrial function in human, macaque, and three other mammalian and avian species. We further map the gene expression of mitochondria-specific pathways in the adult human cortex. The central finding is a consistent dichotomy between anabolic and energy-producing metabolic programs, shared across species and reflected in both the temporal and spatial organization of pathway expression.

In both human and macaque, pathways mainly involved in ATP production including glycolysis, TCA and OXPHOS show a postnatal rise coinciding with the timing of synaptic maturation and peak grey matter volume and cortical thickness (Alldritt et al. 2024, Bethlehem et al. 2022). This finding is also consistent with reported postnatal increases in glycolytic and oxidative phosphorylation proteins in post-mortem proteomic studies (Breen et al. 2018) and with *in vivo* PET imaging of glucose uptake (Chugani 1998, 1999, 2018, Chugani et al. 1987, Goyal et al. 2014, Goyal and Raichle 2018) in human brain development. The OXPHOS pathway particularly shows a peak in the infant and juvenile macaque and decline into adolescence and adulthood. This is in line with our previous findings in human (Pourmajidian et al. 2026) and also *in vivo* PET and nitrous oxide methods measuring of cerebral oxygen consumption during postnatal development in human(Kennedy and Sokoloff 1957, McKenna et al. 2012, Takahashi et al. 1999) and a reduced glucose uptake and blood flow in the brain of aged macaques (Cross et al. 2000, Eberling et al. 1997, Noda et al. 2002). Conversely, the PPP, involved predominantly in tissue biosynthesis shows a pronounced prenatal expression when neural proliferation is highest and declines after birth and progressively in the later ages. This is consistent with previous tracer studies reporting a progressive postnatal decline in pentose phosphate pathway flux in the rat brain, from the early neonatal period to the adult (Baquer et al. 1988, 1977, Brekke et al. 2015, Zubairu et al. 1983). This may also explain the aging-related reduction in aerobic glycolysis, postulated to be a marker of molecular biosynthesis and PPP activity (Goyal et al. 2017).

A parallel pattern emerges within mitochondrial enzymes: pathways of mtDNA maintenance, replication and transcription show greater prenatal expression. Of note is mtDNA nucleoid, a structure that packages the mitochondrial genome, serving as the unit of inheritance during mitochondrial and cell division (Kakudji and Lewis 2024). The greater expression of mtDNA nucleoid genes may therefore reflect the need to expand and partition the mitochondrial pool across rapidly dividing neural progenitor cells during corticogenesis. Conversely, mitochondrial pathways responsible for oxidative, carbohydrate, lipid and amino acid metabolism show a postnatal increase in expression, likely reflecting the postnatal metabolic switch toward mitochondrial energy production (Medini and Mishmar 2025) and the glycolytic-to-oxidative transition observed during neuronal differentiation (Beckervordersandforth et al. 2017, Iwata et al. 2023, Zheng et al. 2016).

Within the mitochondrial lipid metabolism pathways, fatty acid oxidation, cholesterol and phospholipid metabolism peak in childhood, in close alignment with the period of maximum myelination. Of particular note is the decline in genes involved in cardiolipin synthesis across the lifespan. Cardiolipin is a unique phospholipid integral to the formation of tubular cristae within the mitochondrial inner membrane, where it supports electron transport chain supercomplex assembly and the function and efficiency of oxidative phosphorylation. The high content of unsaturated fatty acids in this phospholipid renders it particularly susceptible to peroxidation by the reactive oxygen species generated at the respiratory chain (Ikon and Ryan 2017, Paradies et al. 2014). The age-related decline in cardiolipin expression observed here is consistent with previously reported decreased in cardiolipin content in the aging brain (Paradies et al. 2011, Petrosillo et al. 2008, Sen et al. 2007), likely contributing to the progressive reduction in mitochondrial respiratory capacity with age.

The prenatal-to-postnatal metabolic dichotomy extends beyond the primate brain, with broadly conserved trajectories observed in mouse, rat, and chicken. The postnatal rise in glycolytic and oxidative metabolism is also consistent with measurements of enzyme activity in the rat brain: hexokinase, pyruvate dehydrogenase, citrate synthase, fumarase, and isocitrate dehydrogenase, key enzymes of glycolysis and TCA cycle, all show a postnatal surge in the days after birth (Booth et al. 1980, Clark et al. 1993, Land et al. 1977, Leong and Clark 1984). Furthermore, the activity of the citrate synthase and mitochondrial complexes increase significantly after birth in both synaptosomal and nonsynaptosomal mitochondria isolated from rat brain (Almeida et al. 1995, Bates et al. 1994). The postnatal decline in PPP expression is consistent with previous reports of a progressive decline of this pathway in the rat brain (Baquer et al. 1988, 1977, Brekke et al. 2015, Hakim et al. 1980, Zubairu et al. 1983).

The trajectory of ketone body utilization in mouse and rat aligns with the known reliance of these mammalian species on fat-derived ketone bodies during the suckling period. The expression of monocarboxylic acid transporter in the blood-brain barrier peaks during suckling and declines around weaning as glucose transporter expression rises, reflecting the developmental switch in cerebral fuel from ketone bodies to glucose (Vannucci and Simpson 2003). This is further paralleled by studies on both enzyme activity and cerebral ketone body uptake and oxidation capacity that follow the same temporal pattern (Booth et al. 1980, Clark et al. 1993, Land et al. 1977, Page et al. 1971).

In contrast, the chicken embryo shows a pattern distinct from that of mammals: expression rises during the last week of embryonic development (E14 and E17) and stays elevated around the time of hatching (P0), then subsequently declines in the early postnatal period (P7). This pattern reflects the unique nutritional biology of the avian embryo, which relies on yolk-derived fatty acids as its primary energy source throughout incubation (Speake et al. 1998). In the final week before hatching, yolk lipids are rapidly taken up (Noble and Cocchi 1990, Speake et al. 1998, Yadgary et al. 2010), supporting high rates of ketone body utilization around the time of hatching. The residual yolk sac internalizes into the embryonic body at around E19 and continues to fuel ketone body production into the early postnatal period, before the lipid reserve is gradually exhausted (Speake et al. 1998, Wong and Uni 2021). Blood beta-hydroxybutyrate concentrations also peak at hatching and decline within the first few days post-hatch (Beis 1985). The observed postnatal decline in the expression of ketone body utilization genes thus may reflect the depletion of this lipid reserve, consistent with the reported drop in cerebral 3-hydroxybutyrate uptake in the days following hatching (Linares et al. 1993, Nehlig et al. 1978, Nehlig and Lehr 1981).

Collectively, this convergence across transcriptomic and biochemical evidence implicates these metabolic transitions as fundamental features of vertebrate brain development, and supports the use of neuroimaging transcriptomics in mapping spatial and temporal organization of metabolism in the brain (Colantuoni et al. 2011, Dear et al. 2022, Johnson et al. 2009, Kang et al. 2011, Li et al. 2018, Pletikos et al. 2014, Somel et al. 2010, Vogel et al. 2022).

Finally, the spatial organization of mitochondrial pathways across the adult human cortex mirrors the temporal dichotomy identified across development, suggesting that the anabolic-oxidative distinction is not only a developmental phenomenon but also a persistent feature of cortical organization in the mature brain.

The present study should be interpreted in light of some analytical and dataset limitations. First, the harmonized macaque and human brain transcriptomic dataset has a relatively small sample size, particularly for the childhood stages in the human data. Furthermore, the data used for the additional species comprises only 2–4 samples per time point and is limited to two broad brain divisions (forebrain/hindbrain), making detailed quantitative comparisons difficult across species. We have therefore focused on descriptive comparisons throughout. Second, we relied on orthologous gene mappings to create pathway expression matrices across species. While orthologs are generally assumed to share function, this is not always straightforward. However, core metabolic enzymes are among the most conserved in the genome and here we restricted our analysis to one-to-one orthologs to maximize the likelihood of functional equivalence. This further results in the exclusion of pathway members for which no corresponding orthologs exists, which is especially the case for evolutionarily distant species. Furthermore, mitochondrially encoded subunits are absent from the RNA-sequencing and microarray platforms used here, the trajectories and maps therefore only reflect the nuclear-encoded component of mitochondrial pathway expression. Third, metabolic pathways are highly interconnected and regulated across multiple biological scales; relying on aggregate measures such as mean expression is therefore an oversimplification of the elaborate dynamics governing these pathways. Nonetheless, we have shown that results remain consistent across independent datasets and when using PC1 as an alternative summary measure.

In summary, we leverage existing transcriptomic data to map the lifespan trajectories of brain energy metabolism and mitochondria-specific programs across the mammalian and non-mammalian evolution. The prenatal-to-postnatal metabolic shift, observed as a decline in anabolic PPP expression and a rise in oxidative metabolism, emerges as a conserved feature of the vertebrate brain that closely aligns with the evolving biosynthetic and energetic demands of brain maturation.

## METHODS

### Energy pathway gene sets

Gene sets pertaining to key energy metabolism pathways were curated from Gene Ontology (GO) (Ashburner et al. 2000) and Reactome (Croft et al. 2014) databases using biomaRt version 2.50.3 ((Smedley et al. 2009), https://bioconductor.org/packages/release/bioc/html/biomaRt.html and *GO*.*db* packages (https://bioconductor.org/packages/GO.db/). These pathways are: glycolysis, pentose phosphate pathway (PPP), tricaboxylic acid cycle (TCA), oxidative phosphorylation (OXPHOS), lactate metabolism and transport (lactate) and ketone body utilization. We used the Ensemble human gene annotation database release 112 (Martin et al. 2023) to retrieve gene sets for each pathway ID. Pathway gene sets are provided on our GitHub repository (https://github.com/muhikp/pourmajidian_energy-evolution/). For each pathway, genes annotated in both databases were retained for further analysis to ensure consistent pathway membership.

### Mitocarta3.0 gene sets

Mitocarta3.0 is a public inventory containing gene functional annotations for mitochondrially localized proteins (Calvo et al. 2016, Pagliarini et al. 2008, Rath et al. 2021). Briefly, proteins were identified using in-depth tandem mass spectrometry (MS/MS) of isolated mitochondria from 14 murine tissues, and localization was further assessed by integrating GFP-tagged microscopy, tissue-wide co-expression, and evolutionary evidence. Altogether the authors integrated seven genomics datasets using a naive Bayes classifier trained on known mitochondrial and non-mitochondrial proteins to ensure the accuracy of annotations. Mitocarta3.0 is the most recent release of this dataset that has incorporated manual curation from the literature to update the MitoCarta 2.0 database (Calvo et al. 2016) and further provides functional annotation of 1 136 human genes and a list of 1 140 mouse genes genes into 149 mitochondrial pathways (*MitoPathways* (Rath et al. 2021)). Gene-pathway annotations were downloaded from the MitoCarta website for both human and mouse (https://broadinstitute.org/mitocarta/mitocarta30-inventory-mammalian-mitochondrial-proteins-and-pathways). Pathways are ordered into a three-tier hierarchy. The first level consists of seven major mitochondrial processes: mitochondrial central dogma, protein import, sorting and homeostasis, oxidative phosphorylation, metabolism, small molecule transport, mitochondrial dynamics and surveillance, and signaling. The mitochondrial central dogma pathway refers to mtDNA maintenance and mtRNA metabolism and translation. Out of the 149 annotated MitoPathways, we focused on pathways at the second and third hierarchy levels, which represent progressively finer functional subdivisions within the seven major mitochondrial processes. Among these, we report on pathways that are more relevant to energy metabolism and which showed consistent developmental trajectories across RNA-sequencing and microarray platforms and when using PC1 as an alternative summary measure.

### PsychENCODE human and macaque brain developmental dataset

The PsychENCODE evolution database integrates the human brain transcriptome from Li et al. (2018) with macaque brain transcriptome from Zhu et al. (2018). All experiments using non-human primates were carried out in accordance with a protocol approved by Yale University’s Committee on Animal Research and NIH guidelines. Tissue was collected from macaque specimen that showed no sign of neuropathological alterations, lesions or malformations. The processed data includes normalized Reads Per Kilobase of transcript per Million (RPKM) gene expression values for both human and macaque. RNA-sequencing was performed on bulk tissue from post-mortem specimen and a total of 27 932 genes were probed across 826 samples (*n*_human_ = 460). Human samples come from 36 donors (15 female), across 25 regions (15 cortical) and 26 ages (8 weeks post conception-40 years). The macaque samples come from 26 donors (8 female), across 16 regions (11 cortical) and 16 ages (embryonic day 60-11 years). Annotations of human and macaque orthologous genes were previously performed using the XSAnno pipeline (Zhu et al. 2014). Quality control was done as described in a preceding paper using conditional quantile normalization to account for GC content and sequencing depth and batch effect correction using ComBat (Hansen et al. 2012, Li et al. 2018).

The harmonized developmental transcriptomics for the human and macaque was retrieved from the PsychENCODE website http://evolution.psychencode.org. We carried out a basic clean up of the gene expression matrix. First, we removed duplicate genes. Second, we retained genes that had a log_2_(RPKM) *≥* 1 in at least half of the samples (across both macaque and human). Lastly, expression values were normalized using the upper quartile method (Bullard et al. 2010, Dillies et al. 2013, Miller et al. 2014). Each donor’s data was scaled by their 75th percentile expression value and multiplied by the mean 75th percentile value across all donors. The final matrix contained expression values for 12 509 genes. We then retrieved sample-by-gene expression matrices for each energy pathway using the curated gene sets. Average expression across all genes for each energy pathway was calculated for each sample. Analysis was focused on cortical samples (*n*_human_ = 322, *n*_macaque_ = 257). Cortical regions covered are: medial prefrontal cortex (MFC), orbital prefrontal cortex (OFC), dorsolateral prefrontal cortex (DFC), ventrolateral prefrontal cortex (VFC), primary motor cortex (M1C), primary somatosensory cortex (S1C), inferior posterior parietal cortex (IPC), primary auditory cortex (A1C), superior temporal cortex (STC), inferior temporal cortex (ITC), primary visual cortex (V1C), in addition to four prenatal human regions from motor-somatosensory cortex (MSC), parietal cortex (PC), temporal cortex (TC) and occipital cortex (OC).

Mean energy pathway gene expression was then aggregated into the five age groups (prenate, infant, juvenile, adolescent, adult) by combining all samples within each respective group. Pathway expression was then plotted across age groups. Marker genes for neural progenitor cells (*RPL3, GAPDH, RPL5, RPL7, TMSB15A, PKM, RPL41, TPI1, HNRNPA1, NPM1, LIX1, RPSA, EEF1A1, HMGB2*) neuronal migration (*DCX*, synapse development (*SYP, SYPL1, SYN1*) and myelination (*PLP1, MAG, MBP*) were obtained from the supplementary materials of Kang et al. (2011) and Li et al. (2018). Smoothed curves were produced using the LOESS method against log_10_(age) in post conception days using the *rpy2* package https://rpy2.github.io/doc.html/. For the replication analysis with the BrainSpan microarray dataset, gene expression and sample info matrices were downloaded from https://brainspan.org/static/download.html..Genes were retained if they had a log_2_(expression) *≥* 6 (Kang et al. 2011) and the rest of the analysis was carried out as above.

### Brain developmental gene expression dataset across the vertebrate species

Developmental gene expression data across six mammalian species and chicken were previously published by Cardoso-Moreira et al. (2019) and publicly available through https://apps.kaessmannlab.org/evodevoapp/. Briefly, the authors sampled seven organs including the forebrain and hindbrain from early organogenesis into adulthood. The species sampled are: human, rhesus macaque, mouse, rat, rabbit, opossum and chicken. For this study, we used data from mouse (*Mus musculus*, outbred strain CD-1 RjOrl:SWISS), rat (*Rattus norvegicus*, outbred strain Holtzman SD), and chicken (*Gallus gallus*, red junglefowl). All tissue use was approved by an ERC Ethics Screening panel (ERC Consolidator Grant 615253, OntoTransEvol) and local ethics committees in Lausanne (authorization 504/12) and Heidelberg (authorization S-220/2017).

Brain regions were dissected consistently across the full developmental time series. For mouse, rat and chicken, forebrain samples correspond to cerebral hemispheres (excluding olfactory bulbs). Cerebellum samples correspond to the pre-pontine hindbrain-enriched region up to the developmental equivalent of mouse E15.5, and to the cerebellum proper from E16.5 onwards. RNA was extracted according to the RNeasy protocol from QIAGEN. Read counts were generated using HTSeq (Anders et al. 2015). Count data were normalized using the TMM method in the EdgeR package. Expression tables were provided in RPKM values at https://apps.kaessmannlab.org/evodevoapp/.

First, we removed duplicate genes. Second, expression values were normalized using the upper quartile method (Bullard et al. 2010, Dillies et al. 2013, Miller et al. 2014). Each donor’s data was scaled by their 75th percentile expression value and multiplied by the mean 75th percentile value across all donors. Expression was then log_2_-transformed. Lastly, we retained genes that had an log_2_(RPKM) *≥* 1 in at least 50 percent of the samples. Samples from the earliest developmental stages consist of whole brain without forebrain/cerebellum dissection (Cardoso-Moreira et al. 2019); these were excluded from the present analysis. Developmental trajectories therefore begin at E13 in mouse, E14 in rat, and E10 in chicken.

Ortholog mappings between human and each of the other species were obtained using the *GeneOrthology* package from the Allen Institute https://github.com/AllenInstitute/GeneOrthology NCBI taxonomy IDs were used to retrieve ortholog tables anchored to the human genome (tax ID: 9606) for macaque (*Macaca mulatta*, tax ID: 9544), mouse (*Mus musculus*, tax ID: 10090), rat (*Rattus norvegicus*, tax ID: 10116), and chicken (*Gallus gallus*, tax ID: 9031). Only one-to-one orthologs were retained. We then created sample-by-gene expression matrices for each energy pathway using the mapped ortholog gene sets for each species. Average expression across all genes in each pathway was calculated for each sample. The median pathway expression across samples was then plotted across all ages for each species.

### Microarray gene expression data of the adult human brain

Regional microarray expression data were obtained from 6 post-mortem brains (1 female, ages 24–57, 42.50 ± 13.38; postmortem interval 10–30 hours) provided by the Allen Human Brain Atlas (https://human.brain-map.org). All approvals and consent procedures can be found in the Allen Human Brain Atlas white paper and Hawrylycz et al. (2012). Data were processed using a 400-region volumetric atlas in MNI space as described below. Microarray probes were reannotated using data provided by (Arnatkeviciūtė et al. 2019); probes not matched to a valid Entrez ID were discarded. Next, probes were filtered based on their expression intensity relative to background noise (Quackenbush 2002), such that probes with intensity less than the background in ≥ 50% of samples across donors were discarded, yielding 31 569 probes. When multiple probes indexed the expression of the same gene, we selected and used the probe with the most consistent pattern of regional variation across donors (i.e., differential stability; (Hawrylycz et al. 2015)).

MNI coordinates of tissue samples were updated to those generated via non-linear registration using the Advanced Normalization Tools (ANTs; https://github.com/chrisfilo/alleninf). To increase spatial coverage, tissue samples were mirrored bilaterally across the left and right hemispheres (Romero-Garcia et al. 2018). Samples were assigned to brain regions in the provided atlas if their MNI coordinates were within 2 mm of a given parcel. If a brain region was not assigned a tissue sample based on the above procedure, every voxel in the region was mapped to the nearest tissue sample from the donor in order to generate a dense, interpolated expression map. The average of these expression values was taken across all voxels in the region, weighted by the distance between each voxel and the sample mapped to it, in order to obtain an estimate of the parcellated expression values for the missing region. All tissue samples not assigned to a brain region in the provided atlas were discarded. Inter-subject variation was addressed by normalizing tissue sample expression values across genes using a robust sigmoid function (Fulcher et al. 2013). Normalized expression values were then rescaled to the unit interval (Markello et al. 2021). Gene expression values were then normalized across tissue samples using an identical procedure. Samples assigned to the same brain region were averaged separately for each donor, yielding a regional expression matrix for each donor with 400 rows, corresponding to the cortical regions in the Schaefer-400 parcellation, and 15 633 columns, corresponding to the retained genes. From this initial expression matrix, we retained genes with a differential stability value greater than 0.1 (Burt et al. 2018), yielding expression data for a total of 8 687 genes.

Mitocarta3.0 pathway gene sets were used to extract pathway-specific gene expression matrices. The final number of genes per pathway differs from the original gene sets, as some genes were not present in the AHBA dataset or were excluded based on the above-mentioned quality control criteria applied during preprocessing. For each pathway, expression values were averaged across all genes to yield a mean gene expression pathway map. Brain maps were plotted using the *surfplot* package (Gale et al. 2021, Vos De Wael et al. 2020).

### Spatial auto-correlation preserving nulls

The brain exhibits inherent spatial auto-correlation in both structural and functional measures. Data mapped onto the brain such as gene expression are not independent and identically distributed (i.i.d.), which is a common prerequisite for many statistical tests. Given this spatial autocorrelation, nearby voxels/regions are more likely to have similar values (e.g. similar gene expression) due to both biological and technical (e.g., image processing and smoothing) factors (Markello and Misic 2021). This can lead to inflated statistical values.

To account for this, various spatial permutation tests (spin tests) have been introduced (Alexander-Bloch et al. 2018, Burt et al. 2020, Váša et al. 2018). Spin tests account for the spatial auto-correlation present in brain data by permuting voxels/parcels while maintaining the spatial structure and auto-correlation, therefore providing a more accurate framework for hypothesis testing. In this study, we used the spatial permutation test developed by Váša et al. (2018) implemented in the netneuro-tools package (https://netneurotools.readthedocs.io/). This method uses parcel centroid coordinates to produce rotations and ensure that there are no duplicate reassignments, providing a true null distribution. We refer to the non-parametric p-value calculated using spatial permutation testing as *p*_spin_.

## Data and code availability

All datasets used in this study are publicly available. Scripts and data used to perform the analyses are available at https://github.com/muhikp/pourmajidian_energy-evolution/. Biological databases are openly accessible in *biomaRt* at https://bioconductor.org/packages/release/bioc/html/biomaRt.html. MitoCarta3.0 dataset including the MitoPathways gene sets is available at (https://broadinstitute.org/mitocarta/mitocarta30-inventory-mammalian-mitochondrial-proteins-and-pathways). Processed lifespan brain transcriptomics data is available at http://evolution.psychencode.org/. The lifespan transcriptomics data for the mouse, rat and chicken are available at apps.kaessmannlab.org/evodevoapp/. The Allen Human Brain Atlas is available at https://human.brain-map.org/static/download/. BrainSpan microarray dataset is available at https://brainspan.org/static/download.html. The Schaefer-400 parcellations are openly available at https://github.com/ThomasYeoLab/CBIG/tree/master/stable_projects/brain_parcellation/Schaefer2018_LocalGlobal/Parcellations/ and can also be retrieved using *abagen* at https://abagen.readthedocs.io/.

## Ethics statement

This study uses previously published and publicly available datasets. No new human or animal data were collected as part of this work.

---

**Figure S1.**
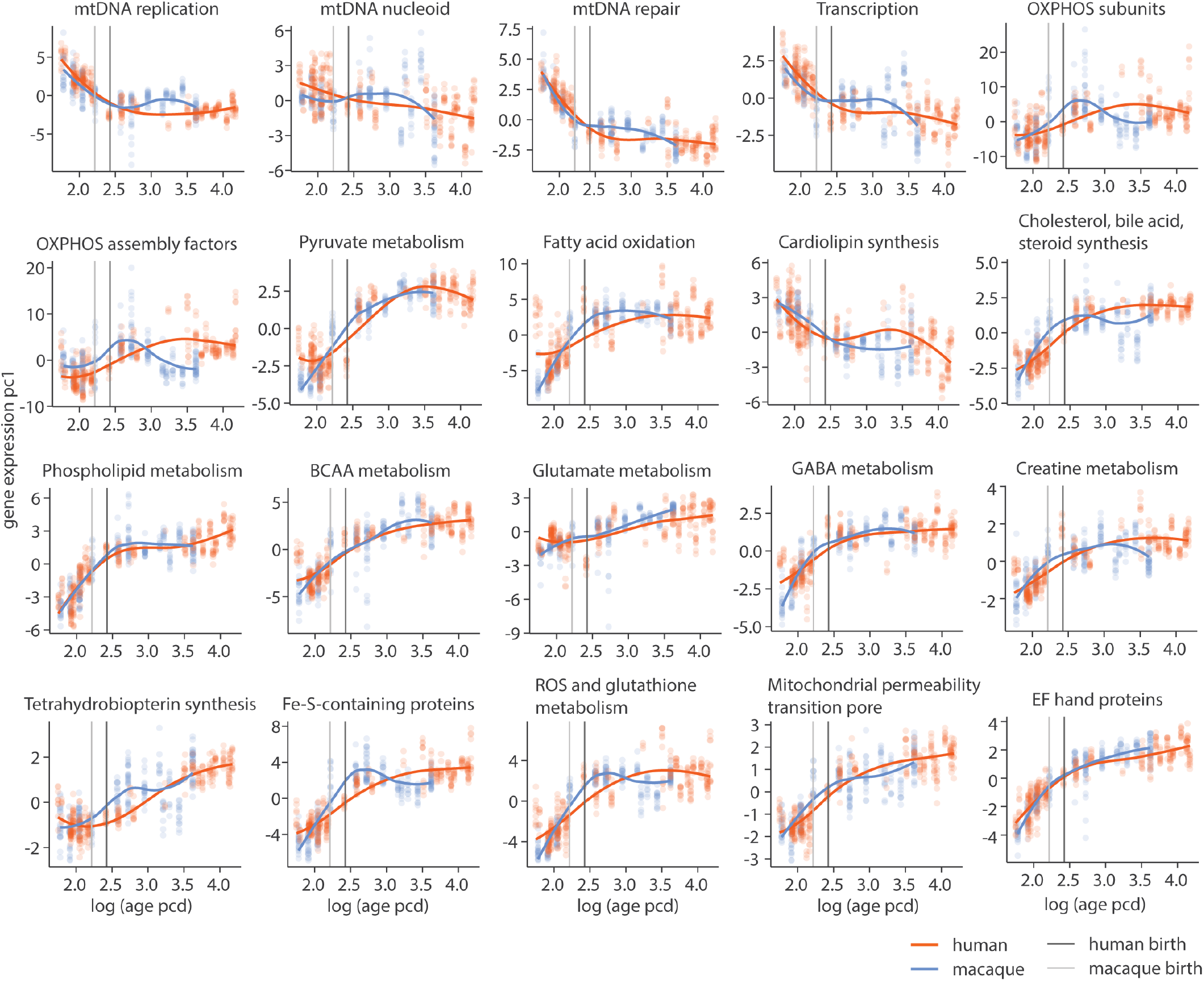
Replication of Mitocarta3.0 pathway trajectories in human and macaque using PC1 of gene expression. As an alternative summary measure to mean expression, the first principal component (PC1) of gene expression was calculated for each pathway gene expression matrix. Smoothed curves were produced using the LOESS method against log_10_-transformed age in post-conception days (PCD). The y-axis represents PC1 scores. Scatter points represent individual samples at each age. Vertical lines indicate macaque and human birth.

**Figure S2.**
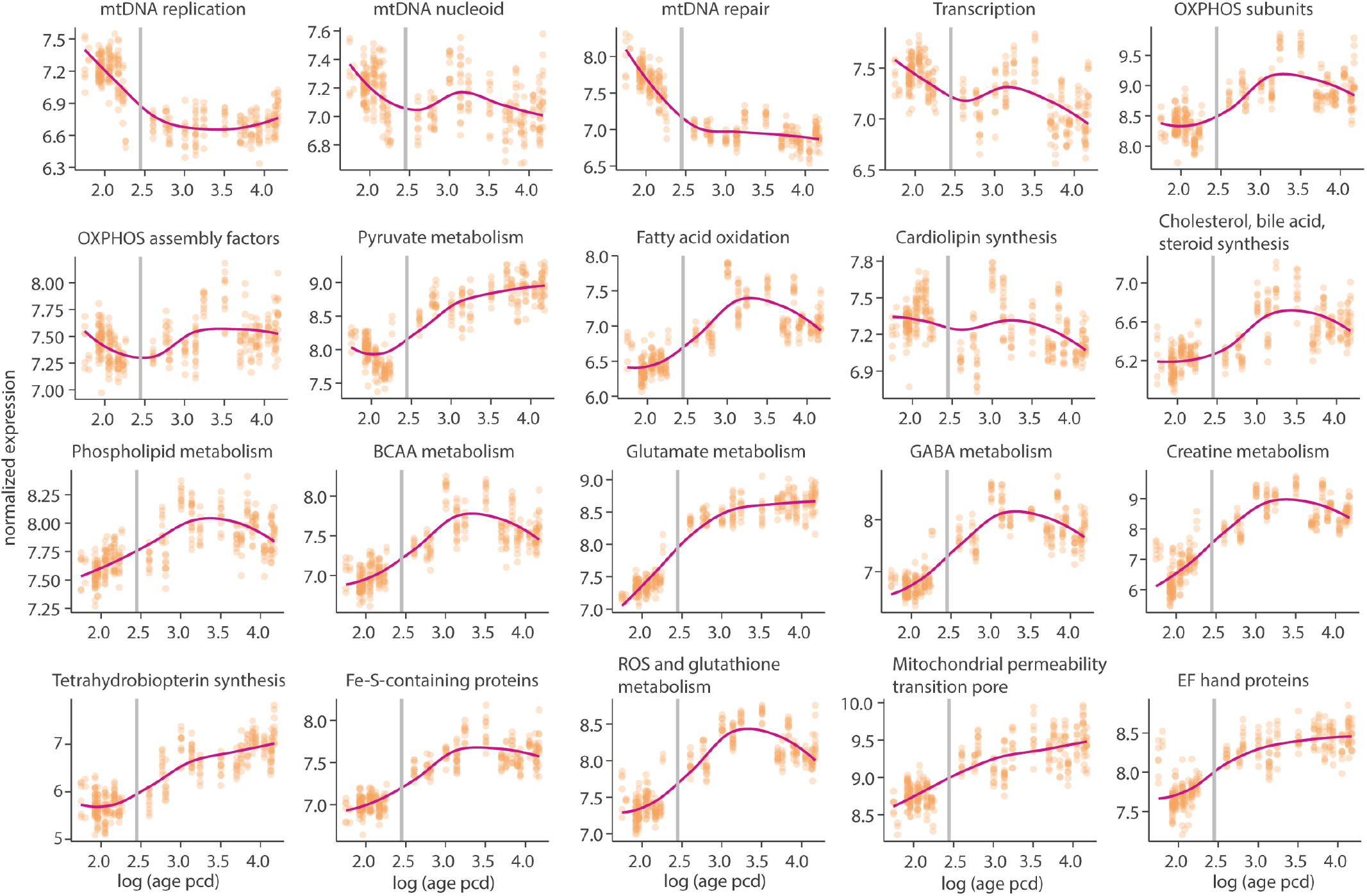
Replication of Mitocarta3.0 pathway trajectories in the human brain using the BrainSpan microarray data. The BrainSpan microarray dataset was used to reproduce pathway sample-by-gene expression matrices (Kang et al. 2011). For each pathway, mean expression was calculated across all available pathway genes. Smoothed curves were produced using the LOESS method against log_10_-transformed age in post-conception days (PCD). The y-axis represents normalized log_2_(RPKM) expression values. Scatter points represent individual samples at each age. vertical line indicates human birth.

**Figure S3.**
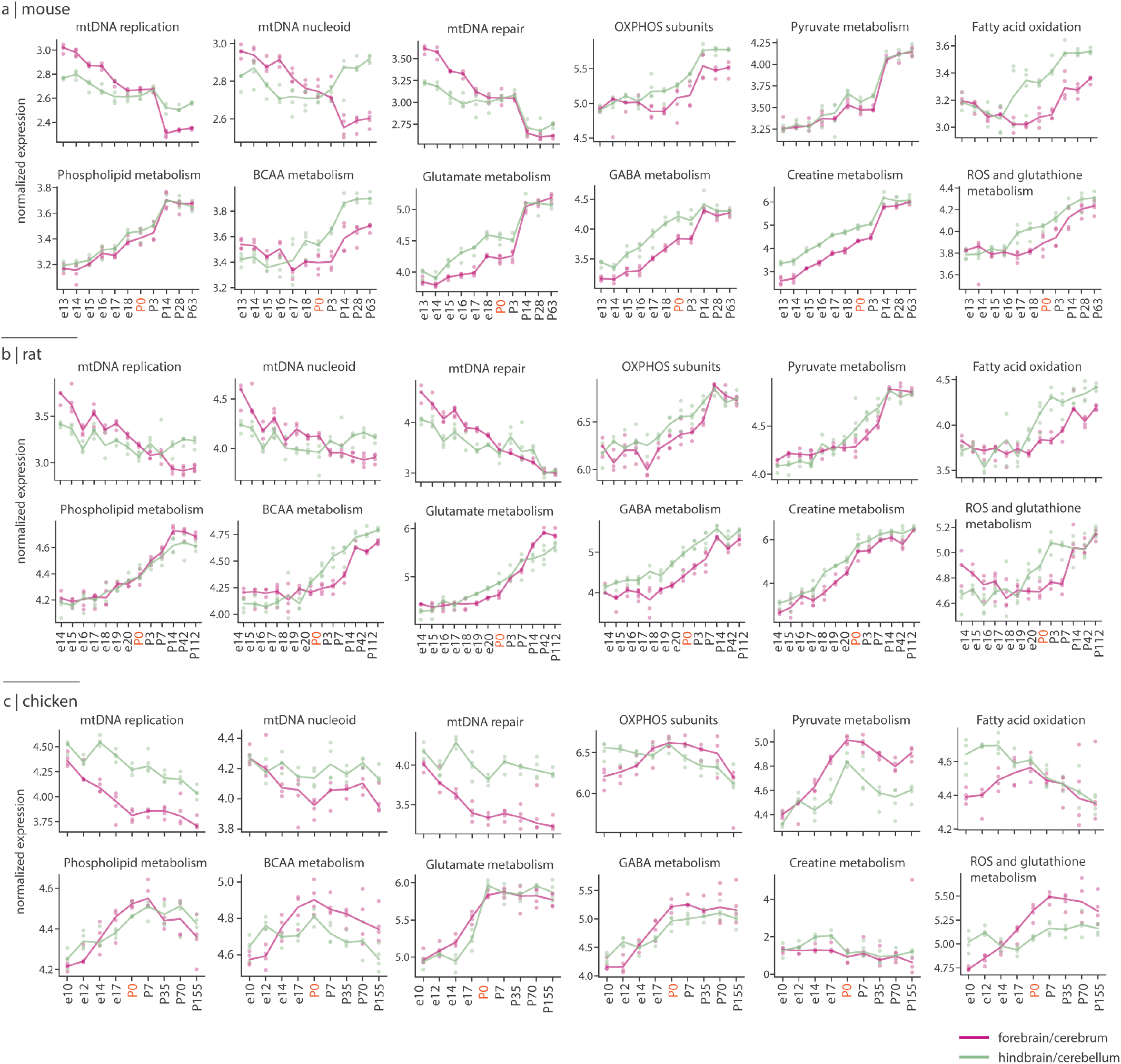
Lifespan trajectories of Mitocarta3.0 pathways across the vertebrate species. Bulk RNA-sequencing data from the forebrain and cerebellum of (a) mouse, (b) rat, and (c) chicken was retrieved from Cardoso-Moreira et al. (2019). For each pathway annotated in Mitocarta3.0 (Rath et al. 2021), sample-by-gene expression matrices were made and mean expression was calculated across all genes for each sample. The median pathway expression across samples at each age was plotted. Pink lines represent forebrain/cerebrum samples; green lines represent hindbrain/cerebellum samples. Scatter points represent individual samples at each age. The y-axis represents normalized log_2_(RPKM) expression values. P0 (red) denotes birth or hatching. MitoCarta3.0 pathway gene sets used in the plots can be found in S1 Data.

**Figure S4.**
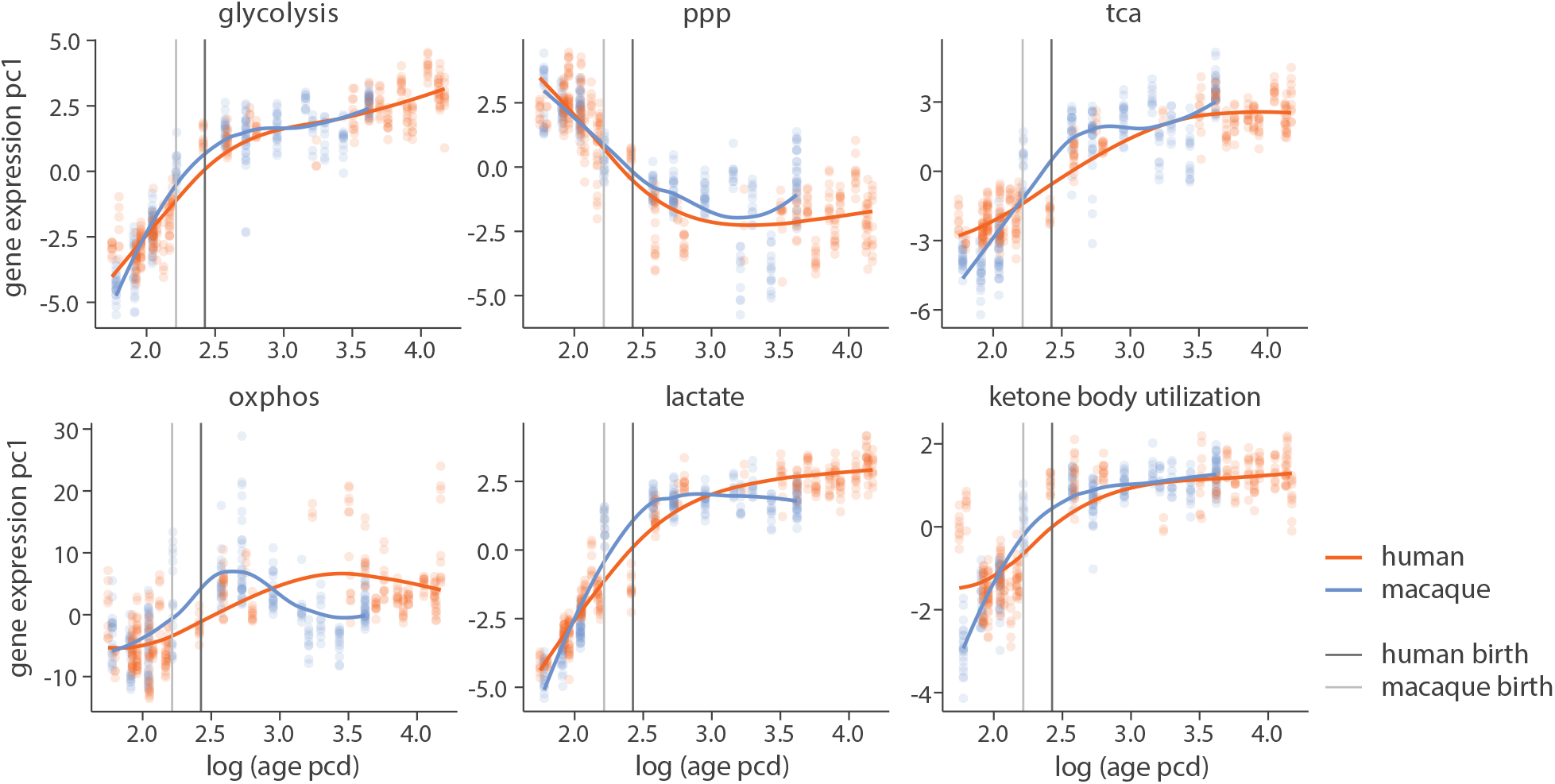
Replication of energy pathway lifespan trajectories in the human and macaque using PC1 of gene expression. As an alternative summary measure to mean expression, the first principal component (PC1) of gene expression was calculated across on the pathwaay sample-by-gene matrices. Smoothed curves were produced using the LOESS method against log_10_-transformed age in post-conception days (PCD). The y-axis represents PC1 scores. Scatter points represent individual samples at each age. Vertical lines indicate macaque and human birth. ppp, pentose phosphate pathway; tca, tricarboxylic acid cycle; oxphos, oxidative phosphorylation; lactate, lactate metabolism and transport.

**Figure S5.**
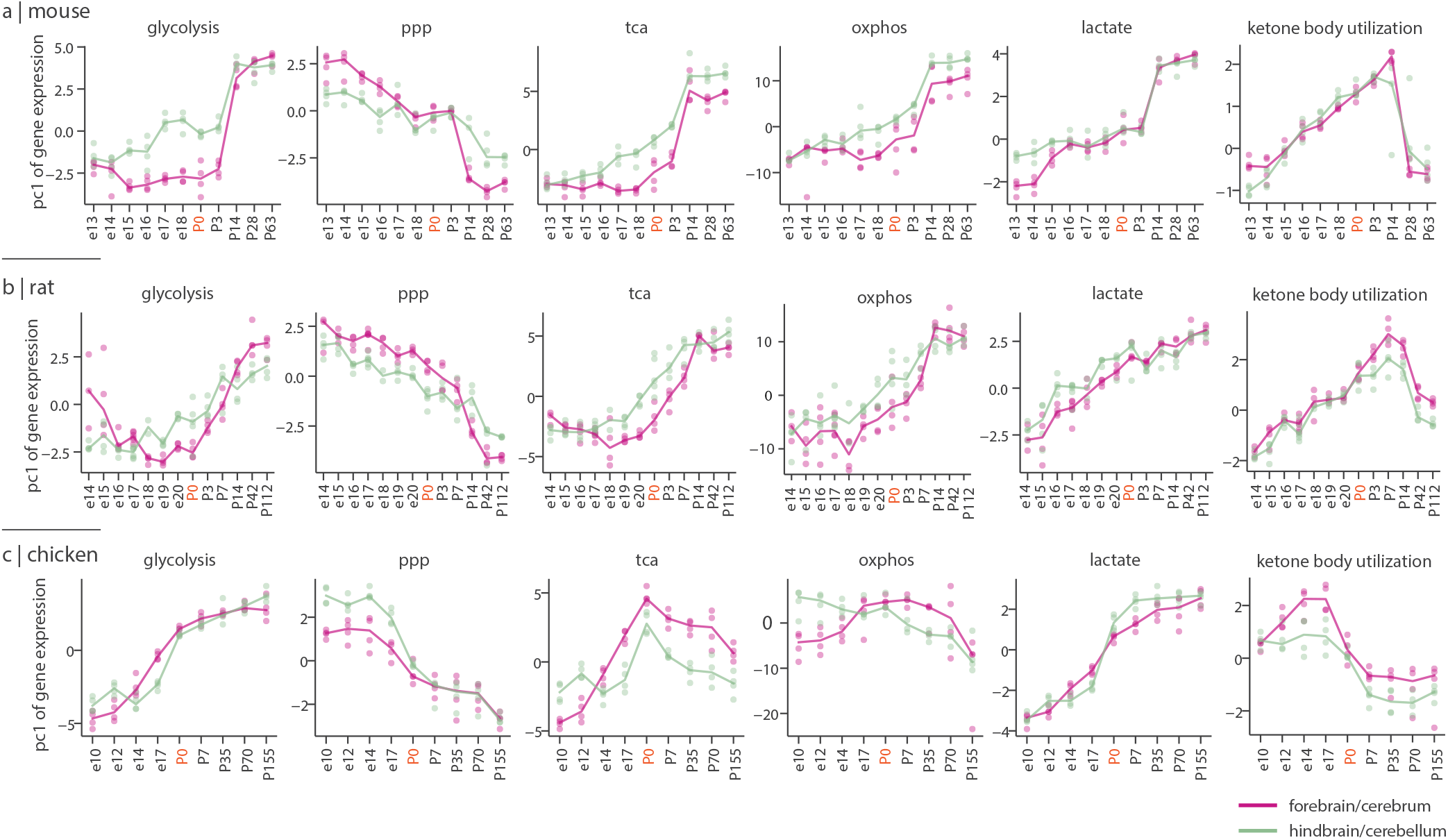
Replication of energy pathway trajectories across vertebrate species using PC1 of gene expression. As an alternative summary measure to mean expression, the first principal component (PC1) of gene expression was calculated for each pathway sample-by-gene expression matrix in the Cardoso-Moreira et al. (2019) dataset. For each species, the median PC1 score of samples at each age is plotted. Pink and green lines represent median gene expression in the forebrain/cerebrum and the hindbrain/cerebellum, respectively. Scatter points represent individual samples at each age. P0 (red) denotes birth or hatching. ppp, pentose phosphate pathway; tca, tricarboxylic acid cycle; oxphos, oxidative phosphorylation; lactate, lactate metabolism and transport.

**Figure S6.**
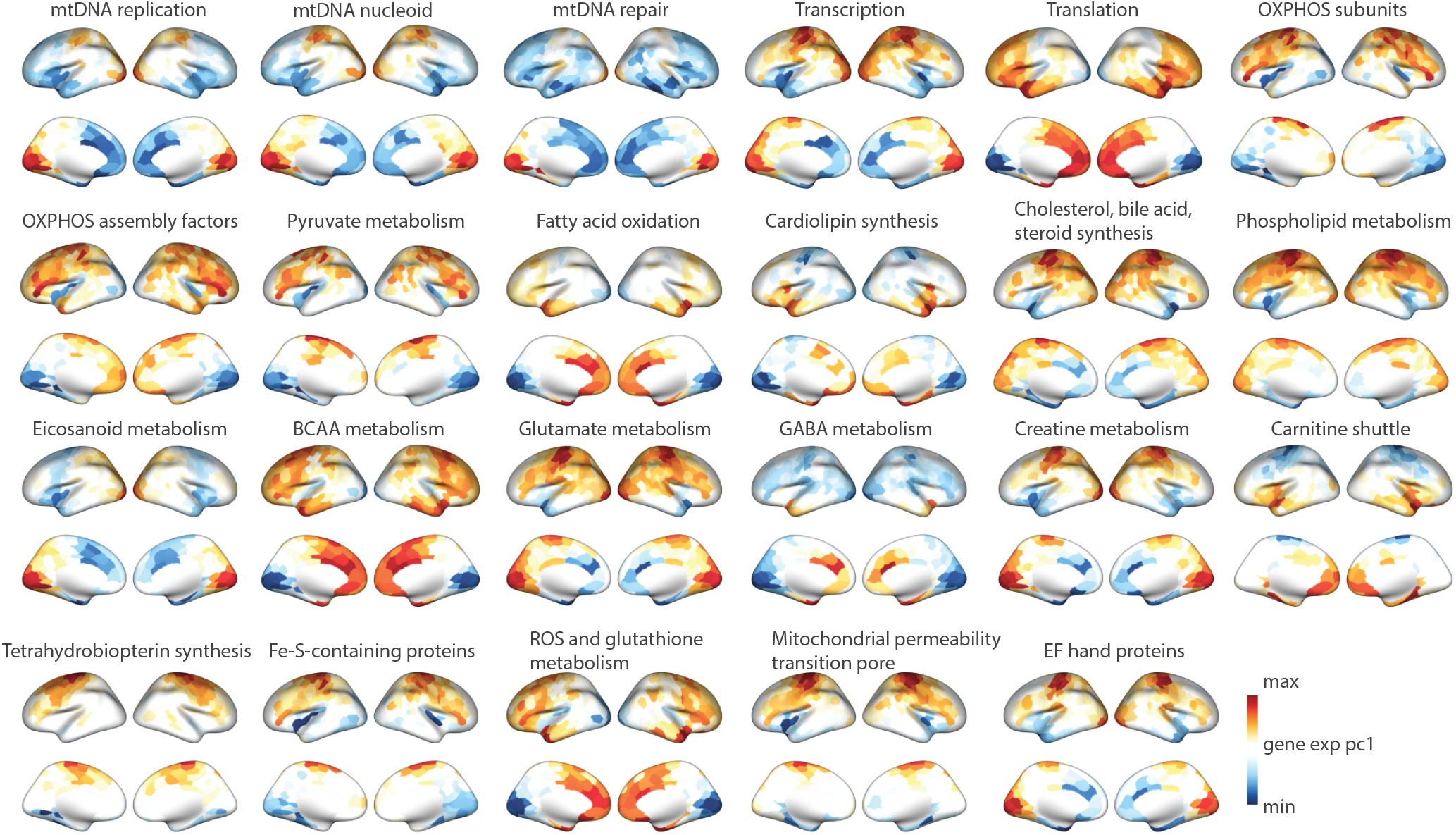
Cortical maps of Mitocarta3.0 pathways using PC1 of gene expression. Pathway-specific gene expression matrices were retrieved from the Allen Human Brain Atlas (Hawrylycz et al. 2012) and parcellated into the Schaefer-400 atlas (Schaefer et al. 2018) using *abagen*. Principle component analysis was done to calculate the first principal component (PC1) of the gene expression matrix for each pathway. Colorbar represents PC1 scores. Data underlying this figure can be found in S1 Data.

**Figure S7.**
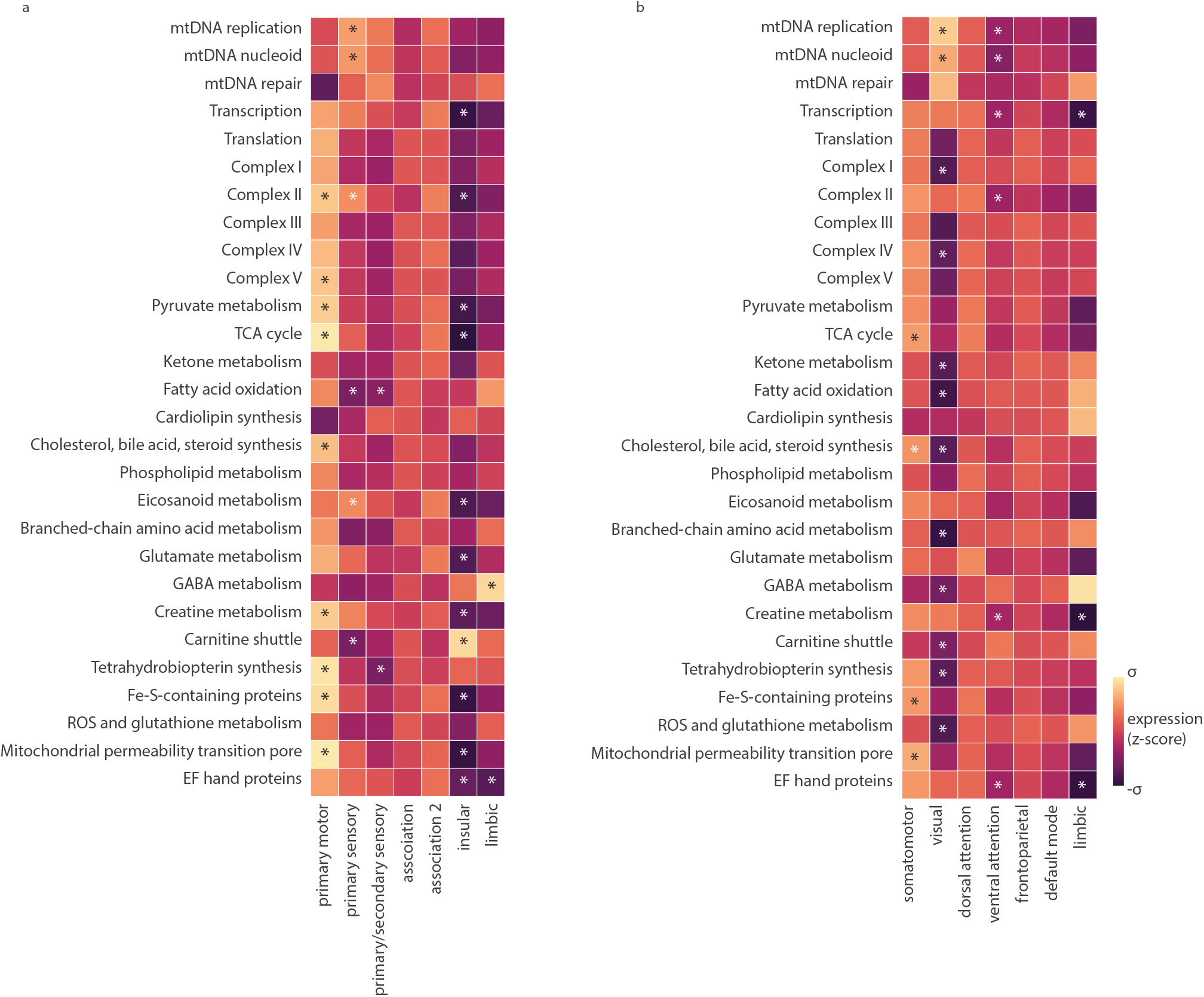
Distribution of MitoCarta3.0 pathway gene expression across structural and functional networks in the cortex. Maps were z-scored across the 400 cortical regions and the average expression of parcels falling into each structural class and functional network was calculated. Distribution of pathway mean expression across (a) seven von Economo-koskinas cytoarchitectonic classes (Economo and Koskinas 1925, Scholtens et al. 2018, Triarhou 2007) and (b) Yeo-Kiernen intrinsic functional network parcellation (Thomas Yeo et al. 2011). Asteriks indicate statistical significance tested against 1 000 spatial-autocorrelation preserving nulls and after FDR correction using the Benjamini-Hochberg method for multiple comparisons (*p*_spin, FDR_ *<* 0.05). Colorbar represents z-scored expression values.

**Figure S8.**
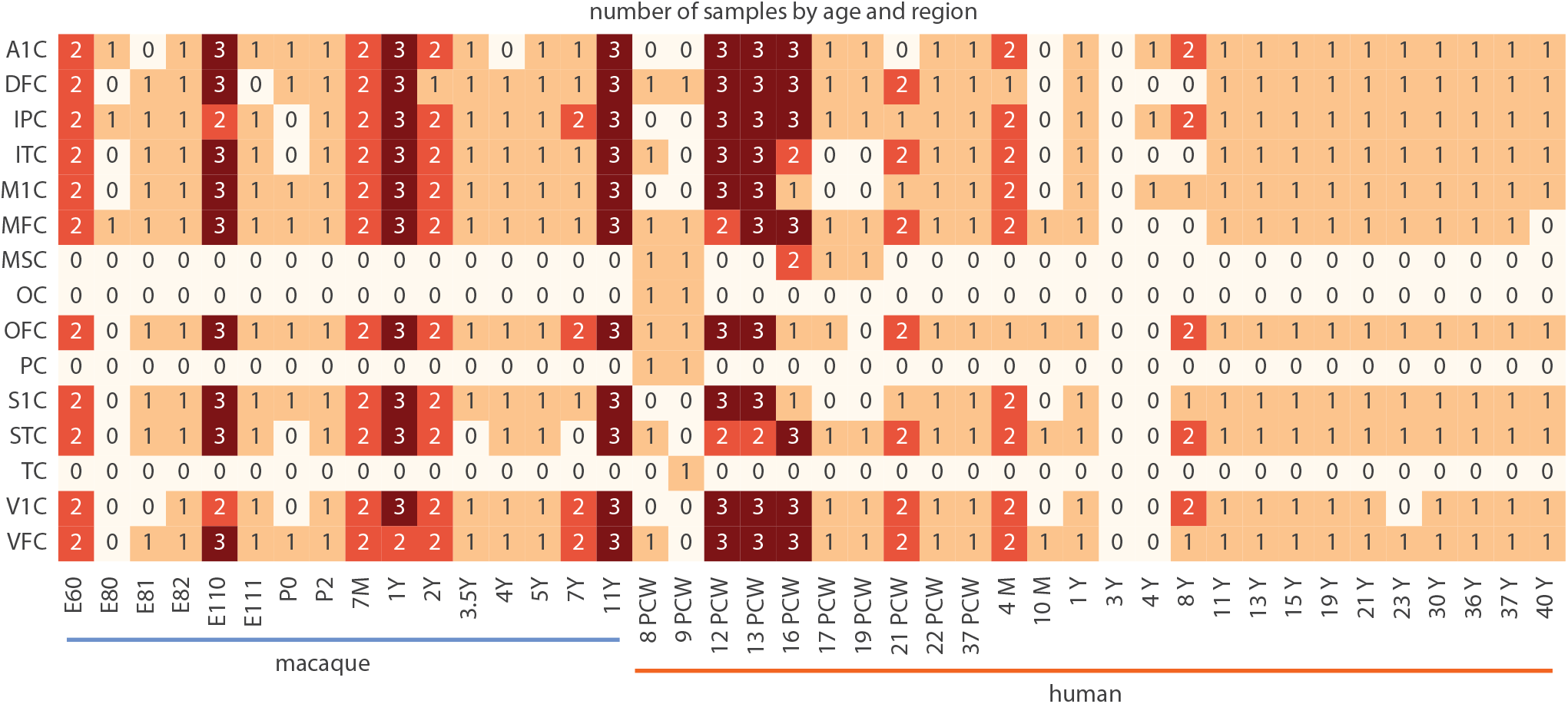
Number of samples at each spatiotemporal point in the PsychENCODE evolution dataset. Sample counts are shown for each brain region and age in the harmonized human and macaque transcriptomic dataset (Li et al. 2018, Zhu et al. 2018). Cortical regions: MFC, medial prefrontal cortex; OFC, orbital prefrontal cortex; DFC, dorsolateral prefrontal cortex; VFC, ventrolateral prefrontal cortex; M1C, primary motor cortex; S1C, primary somatosensory cortex; IPC, inferior posterior parietal cortex; A1C, primary auditory cortex; STC, superior temporal cortex; ITC, inferior temporal cortex; V1C, primary visual cortex. Human prenatal cortical regions: PC, parietal cortex; TC, temporal cortex; OC, and occipital cortex; MSC, motor-somatosensory cortex.

**TABLE S1.**
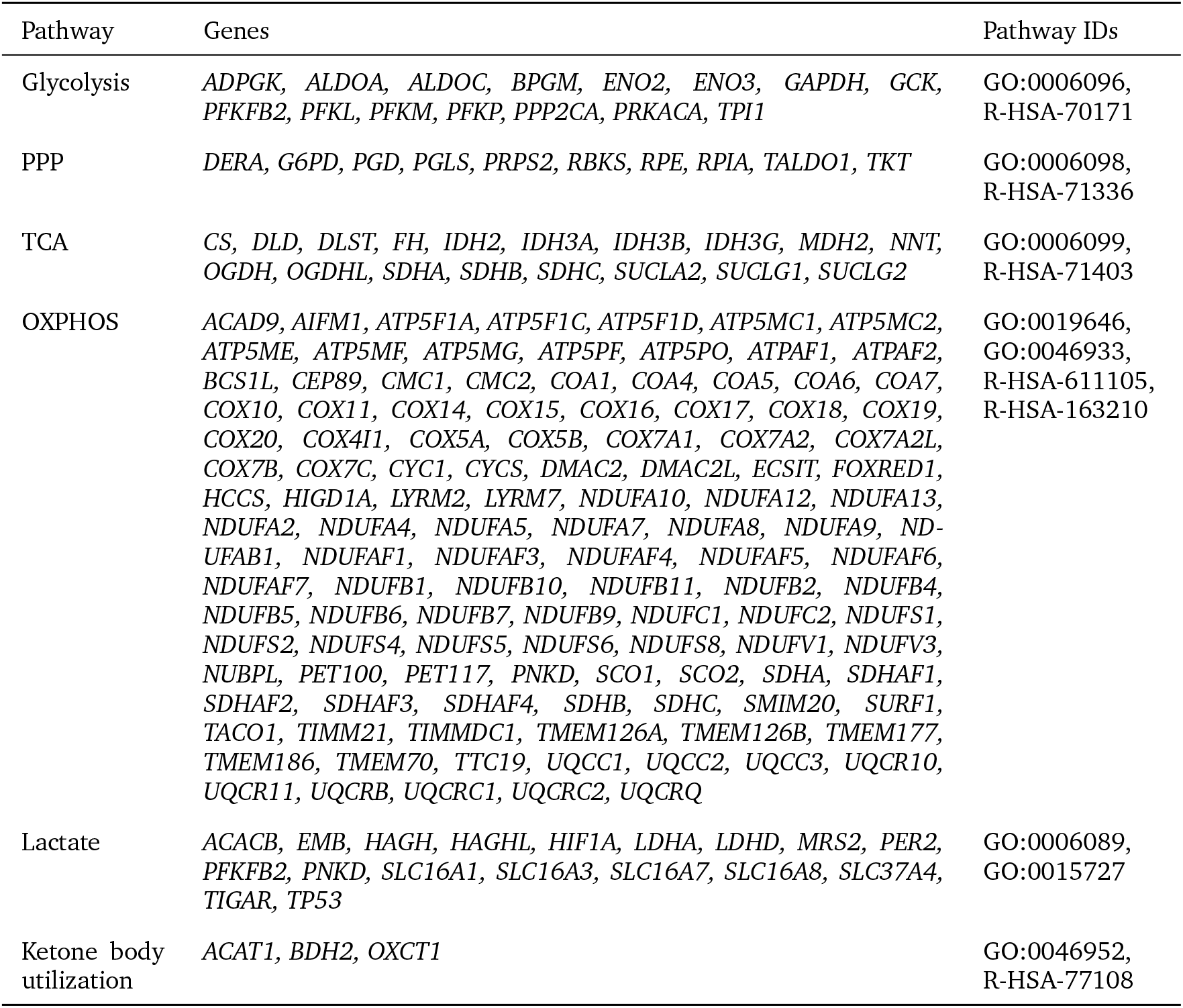
Energy metabolism pathway gene sets. Energy metabolism pathways IDs and gene sets used for the PsychENCODE evolution lifespan analysis (Zhu et al. 2018) in human and macaque. Gene sets were produced based on GO biological processes (Ashburner et al. 2000) and Reactome pathways (Croft et al. 2014) IDs. Genes annotated in both databases and available in the PsychENCODE dataset are listed. ppp, pentose phosphate pathway; tca, tricarboxylic acid cycle; oxphos, oxidative phosphorylation; Lactate, lactate metabolism and transport.

**TABLE S2.** PsychENCODE macaque age groups and sample size. Samples were grouped into five major developmental stages. Last column shows number of cortical samples for each age group used in the lifespan analysis.

| Age group | Age | Num. samples |
| --- | --- | --- |
| Prenate | E60, E80, E81, E82, E110, E111 | 86 |
| Infant | P0, P2, 7M | 40 |
| Juvenile | 1Y, 2Y | 53 |
| Adolescent | 3.5Y, 4Y | 20 |
| Adult | 5Y, 7Y, 11Y | 58 |

S1 Data. **Supplementary tables containing the final gene sets used for the main energy metabolism and MitoCarta3.0 pathway analyses, and numerical data for Figure 5**.

